# Lifetime of actin-dependent protein nanoclusters

**DOI:** 10.1101/2022.01.06.475299

**Authors:** Sumantra Sarkar, Debanjan Goswami

## Abstract

Protein nanoclusters (PNCs) are dynamic collections of a few proteins that spatially organize in nanometer length clusters. PNCs are one of the principal forms of spatial organization of membrane proteins and they have been shown or hypothesized to be important in various cellular processes, including cell signaling. PNCs show remarkable diversity in size, shape, and lifetime. In particular, the lifetime of PNCs can vary over a wide range of timescales. The diversity in size and shape can be explained by the interaction of the clustering proteins with the actin cytoskeleton or the lipid membrane, but very little is known about the processes that determine the lifetime of the nanoclusters. In this paper, using mathematical modelling of the cluster dynamics, we model the biophysical processes that determine the lifetime of actin-dependent PNCs. In particular, we investigated the role of actin aster fragmentation, which had been suggested to be a key determinant of the PNC lifetime, and found that it is important only for a small class of PNCs. A simple extension of our model allowed us to investigate the kinetics of protein-ligand interaction near PNCs. We found an anomalous increase in the lifetime of ligands near PNCs, which agrees remarkably well with experimental data on RAS-RAF kinetics. In particular, analysis of the RAS-RAF data through our model provides falsifiable predictions and novel hypotheses that will not only shed light on the role of RAS-RAF kinetics in various cancers, but also will be useful in studying membrane protein clustering in general.

**Significance:** Spatial organization of biomolecules shapes the behavior of a cell. It is particularly important during cell-signaling, where transient, dynamic organization of the biomolecules helps cells process signals and respond to them. Nanoclusters of peripheral membrane proteins, such as KRAS, a specific form of dynamic organization of biomolecules, play a critical part in the modulation of cell signals that control various cellular behaviors including cell growth, proliferation, and differentiation. Although we have made significant progress in understanding the structure, size, and origin of the nanoclusters, very little is known about the biophysical processes that control their lifetime. In this paper, we present a mathematical framework that provides quantitative insights into these processes and explains how oncogenic mutations in KRAS may lead to cancers.

## Introduction

Protein nanoclusters (PNCs) are dynamic collections of a small number of proteins that spatially organize in nanometer length clusters [1–9]. PNCs are one of the principal forms of spatial organization of membrane proteins and they have been shown or hypothesized to be important in various cellular processes. In particular, it has been postulated that PNCs can digitize noisy analog signals that improve the signal-to-noise ratio of the signals received by a cell [10–14]. Furthermore, it has been shown theoretically that PNCs can drastically improve the reaction rates of double-modification networks by allowing rapid multiple rebinding of an enzyme and its substrate [15]. Importantly, the presence or absence of PNCs have measurable impact on the cell physiology and cellular behavior. For example, in mast cells, which control the response to allergic reactions, proliferation of the Fc*ε*R receptor clusters has been linked with the degranulation of the cells and strong allergic response [16]. In another example, the formation of glycosylphosphatidylinositol-anchored protein (GPI-AP) clusters have been shown to be important in mechano-sensing and cell-spreading [17]. Therefore, understanding the dynamics of formation, growth, function, lifetime, and disintegration, of PNCs is of paramount importance in our pursuit to understand and control cell signaling and the various diseases that it engenders.

Depending on the specific function and the cell-type, the composition, size, shape, and the lifetime of the PNCs vary a lot. They can be homogeneous in composition, e.g., Kirsten rat sarcoma virus (KRAS) protein nanoclusters [3], or heterogeneous, such as in focal adhesion clusters [17,18]. The shape can be isotropic, as of KRAS [4], or anisotropic, as of Harvey RAS (HRAS) PNCs [19]. The number of proteins in a PNC (cluster size) also varies over quite a range. For example, KRAS nanoclusters typically contain 3-8 proteins [3,4], whereas Fc receptors can form clusters of 20-30 proteins [16]. Similarly, the cluster radius can also vary from 20-200 nm [9]. Finally, the lifetime and the stability of the clusters can also vary over a broad range: KRAS clusters, which are *peripheral* membrane protein clusters, are transient and survive for around 0.1-1 s [3,4], whereas Fc receptor clusters, which are *integral* membrane protein clusters, are stable and can survive the entire duration of the experiments (minutes) [16]. Protein-protein interactions and protein-lipid interactions play a key role in determining the size, shape, composition, and stability of the PNCs specific to a biological process. Therefore, to understand the dynamics of PNCs in specific processes and cell types, several studies have investigated the underlying protein-protein and protein-lipid interactions [4,9]. Despite the diversity of the PNCs and the underlying systems, these studies have shown that the formation of the PNCs can be categorized into actin-dependent and actin-independent groups [9], which provides a general framework for studying the dynamics of the PNCs. In this paper, we focus on the dynamics of actin-dependent peripheral membrane protein clusters.

The formation of the actin-dependent peripheral membrane protein clusters (PNCs henceforth) happens through a set of biomolecular processes that can be abstracted into a simple physical model, first developed to understand actin-dependent clustering of GPI-AP [20]. In this model, described in Fig.1, the formation of actin asters aids the PNC formation. In particular, it is assumed that the protein adsorbs on the actin fiber with a rate *k_on_*, advected to the aster center, and desorbs with a rate *k_off_*, which leads to the formation of a dynamic PNC. Importantly, the protein absorption kinetics does not impact the actin self-organization dynamics in any ways, but the actin asters do impact the PNC lifetime through the fragmentation of the asters. One corollary of these assumptions is that the aster lifetime solely determines the PNC lifetime. However, experimental observations do not corroborate this statement. In particular, recent experiments have revealed that asters survive for 10-500 seconds in *in vivo* conditions and their typical fragmentation times are around 20s [21,22]. In contrast, many actin-dependent clusters, such as KRAS clusters, survive for only 0.1-1 s [4], which suggests that the lifetime of a PNC is determined by multiple physicochemical processes besides the aster fragmentation. Indeed, a prior work has suggested that stochastic protein absorption kinetics can be one such mechanism [23]. However, it remains unclear whether the PNC lifetime is always determined by the stochastic growth kinetics or whether it is determined by both actin fragmentation and adsorption kinetics, depending on the specific situation. In this paper, we propose a model of PNC growth kinetics that, to the best of our knowledge, allows us to answer this question for the first time. Our model shows that although the formation of actin asters is necessary for the formation of actin-dependent clusters, under most biological conditions, they do not determine the lifetime of PNCs.

**Figure 1.**
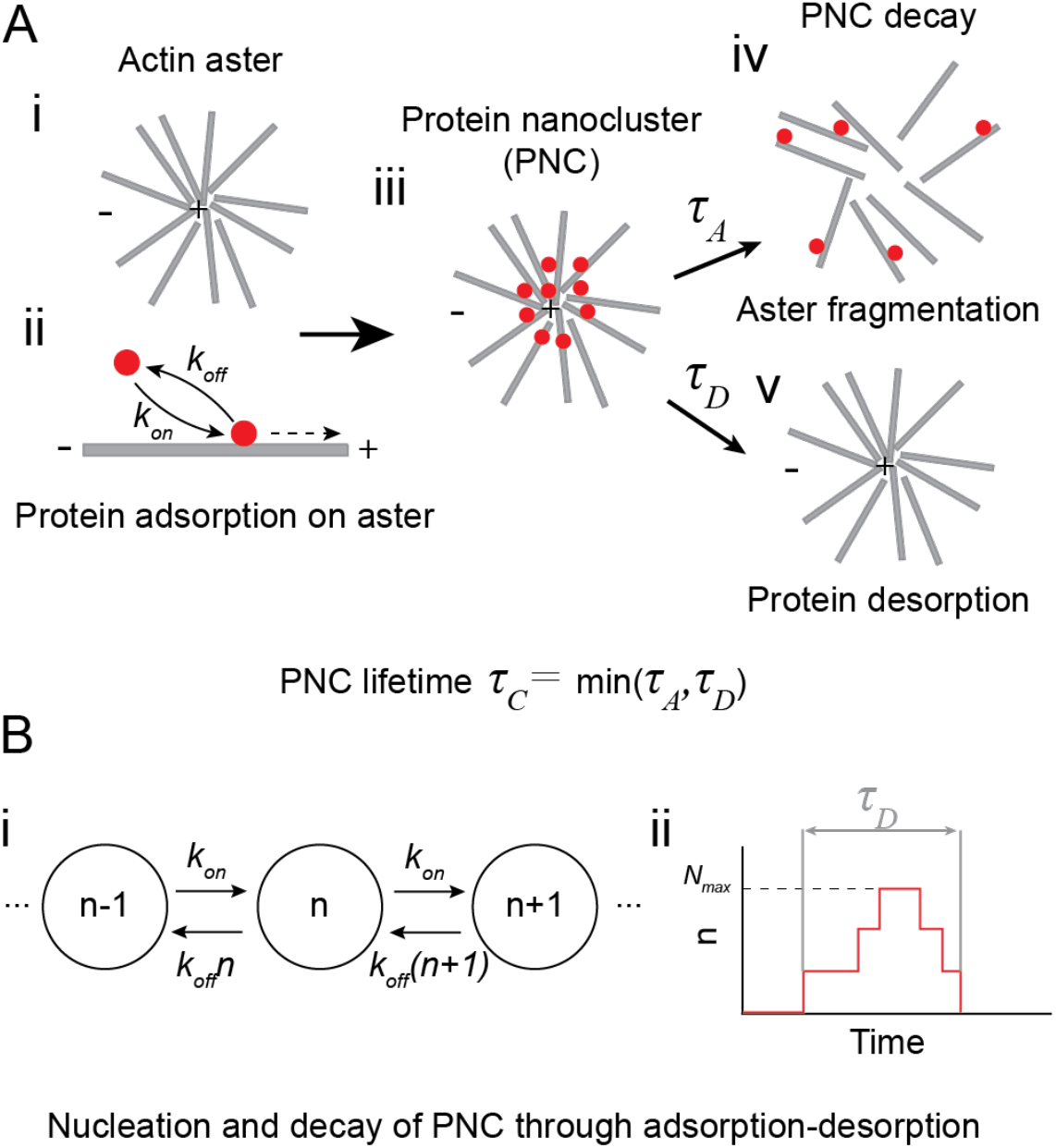
The aster-driven nucleation of protein nanoclusters: (A) (i) Dynamic cortical actin fibers forms asters on the cell membrane. (ii) Peripheral proteins bind (adsorbs) to the dynamic actin fiber with rate *k_on_* and are advected to the “+” end of the fiber. Once bound, it dissociates (desorbs) from the membrane with rate *k_off_*. (iii) Proteins adsorbed on the membrane and advected to the plus end of the aster form a protein nanocluster. The protein nanocluster disintegrates either through (iv) the fragmentation of the aster with timescale *τ_A_* or (v) through the desorption of all the proteins with timescale *τ_D_*. The lifetime of the cluster is the minimum of these two times. (B) The cluster formation model. (i) The cluster size *n* grows with rate *k_on_* and decays with rate *k_off_* × *n*. (ii) The cluster grows from *n =* 0 to a maximum size *n = N_max_* and then decays back to *n =* 0, after time *τ_D_*. We define *N_max_* as the cluster size.

As an application of the general model presented in this paper, we also investigate the formation of heterogeneous protein clusters, such as the KRAS-C-RAF clusters, and their lifetime through a simple extension of this model. KRAS4B (RAS henceforth) is a peripheral membrane protein in human cells which interacts with the kinase C-RAF (RAF henceforth) to control cellular growth, differentiation, and proliferation [24–26]. Mutated RAS and RAF is the underlying cause of 30% of all known cancers [27,28]. In this paper, through the application of our general model we have shown how oncogenic mutations in RAS lead to cancers. The insights gained from our quantitative predictions may be useful in developing therapeutics against these cancers.

## Results

### Determinants of protein nanocluster lifetime

To understand the processes that determine PNC lifetime, we used a simplified version of the model proposed by Gowrishankar et.al. [20] In particular, we assume that the aster formation happens at a timescale much faster than the PNC lifetime, such that the aster has already reached its steady-state structure when the first protein adsorbs on the aster. This assumption decouples the transient PNC kinetics from the transient aster dynamics, and we can incorporate the contribution of the aster on the PNC kinetics simply through the local actin concentration *C*. Under this assumption, the PNC lifetime is determined by two processes: (a) the stochastic growth and decay of the PNC through protein adsorption and desorption, which gives a desorption time *τ_D_*, and (b) aster fragmentation with mean fragmentation time *τ_A_* (Fig 1A). The cluster lifetime *τ_c_* is given by the shorter of these two times. That is:

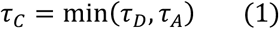

To better understand the origin of *τ_D_*, consider the model shown in Fig. 1B(i). The proteins adsorb onto the aster at a propensity *k_on_C*, which increases the size of the cluster by one protein. The size of the cluster decreases through the desorption process, which, for a cluster of size *n*, happens at a propensity *k_off_n*. We also assume that all asters are identical, such that they have identical actin concentration *C*, which allows us to absorb *C* in *k_on_*, such that the propensity of growth is *k_on_*. We relax this assumption later to include variable *C* also. Due to the small number of proteins, the growth and decay of the cluster is a stochastic process in which the size of the cluster grows from *n* = 0 to a maximum size *N_max_* and then ultimately decays back to *n* = 0. *We define the total time between these two n = 0 states as τ_D_ and the cluster size is given by N_max_* (Fig 1B(ii)).

To understand the relative contribution of the stochastic growth and the aster fragmentation on the lifetime of a PNC, we measured the distribution of *τ_D_*, *P_D_*(*τ_D_*), by varying *k_off_* and the duty ratio 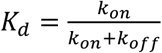, which measures the fraction of time a single protein remains bound to the actin. Following experimental observations [21,22], we assumed that the aster fragmentation time, *τ_A_*, is exponentially distributed with ⟨*τ_A_*⟩ = 20 *s*. Given *P_D_*(*τ_D_*) and *P_A_*(*τ_A_*), the cluster lifetime distribution is given by:

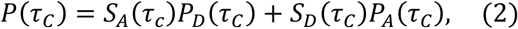

where *S_A_* and *S_D_* are the survival probability functions of *P_A_* and *P_D_* (see SI). This expression simply states that the cluster lifetime *τ_c_* is determined by *P_D_*, if the aster survives until time *τ_c_*, else it is determined by *P_A_*. In Fig. 2A, we show some example distributions. When both *k_off_* is small and *K_d_* is large, a protein adsorbs at a rate much higher than it desorbs and once adsorbed takes a long time to desorb. We find that, in these situations, the cluster lifetime is determined by the aster fragmentation time (Fig 2A (i), (iii)). In contrast, for all other situations, *P*(*τ_c_*) is determined by *P*(*τ_D_*) (Fig 2A (ii), (iv)). We can make this observation more quantitative by measuring the overlap between *P*(*τ_c_*) and *P_D_*(*τ_c_*) or *P_A_*(*τ_c_*). In Fig. 2B, we show the overlap between *P*(*τ_c_*) and *P_A_*(*τ_c_*), which shows that protein desorption determines the cluster lifetime for a large set of parameters. Only when the adsorption-desorption process is extremely slow and the protein strongly adsorbs to the actin fiber, then the lifetime is determined by the aster fragmentation time. This observation shows that while actin aster is necessary for the formation of actin-dependent PNCs, in cellular conditions, its fragmentation may rarely determine the PNC lifetime. A simple testable prediction from this model is that, as the aster fragmentation time shortens, e.g., through the application of Latrunculin [2], there will be a sharp transition in the cluster lifetime distribution.

**Figure 2.**
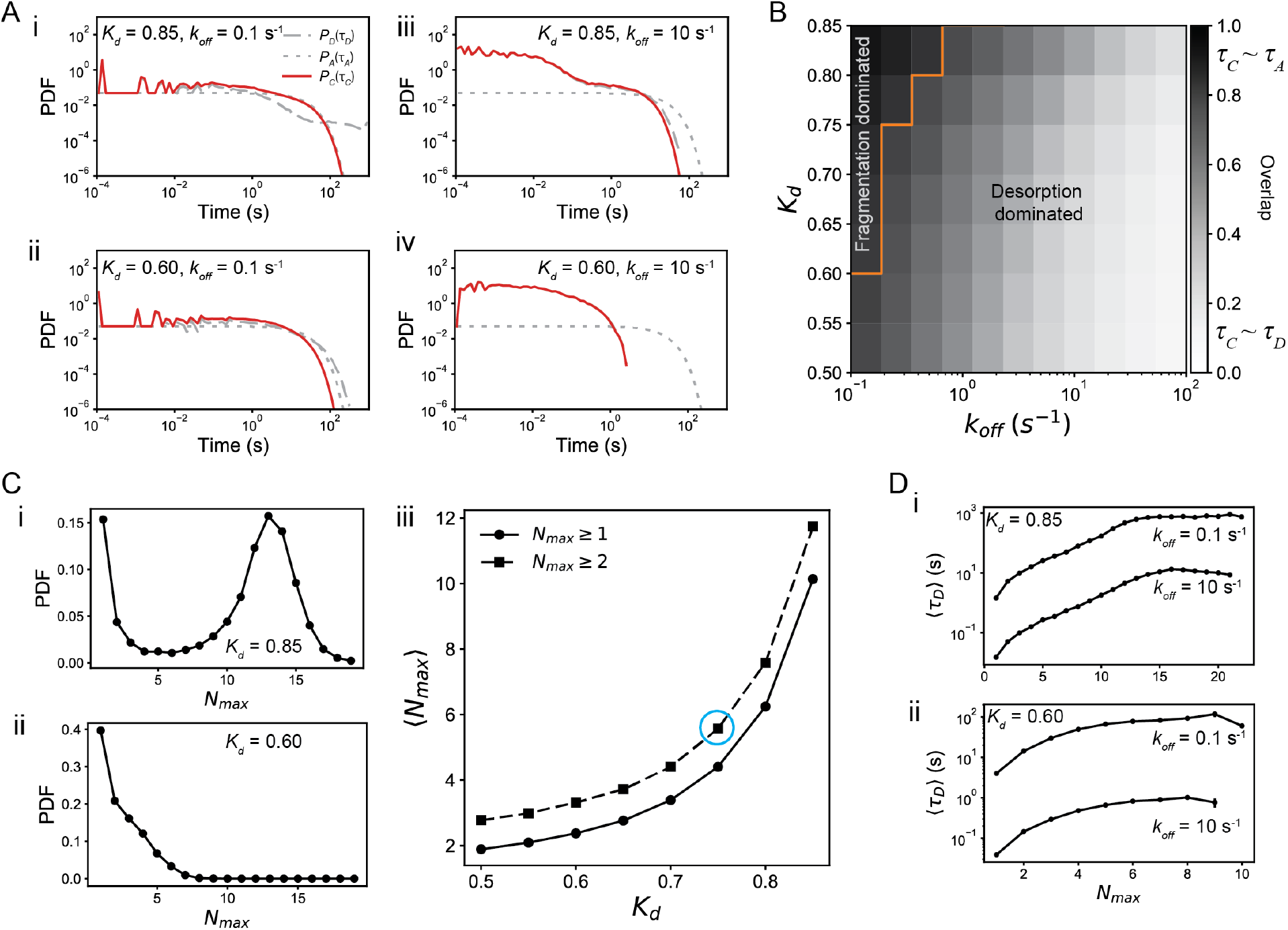
Protein nanocluster lifetime distribution. (A) Cluster lifetime (solid red line) and desorption time (gray dashed line) for different values of *K_d_* and *k_off_*. For all simulations, we assume that the aster lifetime is exponentially distributed with mean lifetime of 20 s (gray dotted line). The cluster lifetime is more similar to the aster lifetime when *K_d_* is high and *k_off_* is low (i,ii), because for such parameters, the cluster is extremely stable and cluster decay happens because of aster fragmentation. Similarly, when *K_d_* is low or *k_off_* is high (iii,iv), the cluster lifetime is more similar to the desorption timescale, because the clusters are much less stable. The similarity of *P*(*τ_C_*) with *P*(*τ_A_*) or *P*(*τ_D_*) can be measured by finding the overlap between these distributions. For example, in (B) we show the overlap between *P*(*τ_C_*) and *P*(*τ_A_*), which consolidates our observation that cluster lifetime is similar to aster lifetime when *K_d_* is high and *k_off_* is low. The orange line marks the boundary where the overlap of *P*(*τ_c_*) with *P*(*τ_A_*) is higher than that with *P*(*τ_D_*). (C) The distribution of cluster size, *N_max_*, depends on *K_d_*, but not on *k_off_*. The distribution for (i) *K_d_* = 0.85 and (ii) *K_d_* = 0.60 shows that as *K_d_* increases, the distribution develops a peak at *N_max_* > 1, which leads to the nonlinear increase of the average ⟨*N_max_*⟩ with *K_d_*. (D) This change in the cluster size distribution drastically impacts the average desorption time ⟨*τ_D_*⟩. When (i) *K_d_* = 0.6, the average lifetime grows beyond the average aster lifetime, ⟨*τ_A_*⟩ only when *k_off_* is small, but (ii) for *K_d_* = 0.85, the average lifetime grows rapidly beyond ⟨*τ_A_*⟩. These two results reaffirm the location of the boundary in (B). The blue circle in C(iii) indicates the value of *K_d_* that is most similar to experimentally observed RAS cluster size distribution. 10^4^ independent realizations of the model were used for these results.

### Dependence of desorption time on the cluster size

The cluster size, i.e., *N_max_*, does not depend on the desorption rate *k_off_* and depends only on the duty ratio *K_d_*. As *K_d_* increases, the cluster size distribution transitions from a unimodal distribution to a bimodal distribution, implying that larger clusters become more prevalent at higher *K_d_* values. Indeed, the mean size of the clusters increase nonlinearly with *K_d_*, being close to 1 for *K_d_* ≈ 0.5. The average desorption time, ⟨*τ_D_*⟩ has a nontrivial dependence on the cluster size and *k_off_*. In particular, ⟨*τ_D_*⟩ vs *N_max_* shows three different regimes. When *N_max_* is small, ⟨*τ_D_*⟩ increases subexponentially with *N_max_*, followed by exponential increase, and saturation to a maximum value that depends on *K_d_* and *k_off_*. For example, for *K_d_* = 0.6, ⟨*τ_D_*⟩ saturates to a value that is much smaller than the simulation ceiling (1000 s), whereas for *K_d_* = 0.85 and *k_off_* = 0.1*s*^−1^, the saturation happens at the simulation limit, implying that the ⟨*τ_D_*⟩ values are much longer than the simulation ceiling. This observation also clarifies why we observe fragmentation dominated cluster lifetime only when *k_off_* is small and/or *K_d_* is large. As our results show, only in this limit the cluster sizes are large enough such that *τ_D_* is much longer than *τ_A_*.

### Ligand-protein interaction on protein nanoclusters

One recurrent feature of cell signaling systems is that cytosolic or extracellular ligands are recruited to a membrane PNC in response to a signal [29]. For example, the effector protein RAF is recruited to clustered RAS, which starts a MAPK signaling pathway for cell growth and proliferation [24,25]. Therefore, it is important to understand how the clustering of the membrane proteins influence and modulates the ligand dynamics. Investigation of this problem is particularly exciting when the membrane protein forms desorption dominated PNCs, because in such a situation the ligands form a transient cluster whose growth kinetics is intimately coupled to the growth kinetics of the underlying PNC. For fragmentation dominated PNCs, because of their long lifetime, the ligand growth kinetics is very similar to the PNC kinetics. Because of this reason, in this paper, we are only going to focus on the ligand-protein interaction on desorption dominated PNCs.

To model the ligand-protein interaction, we extend our model of PNC formation by including an additional molecular species (blue particles in Fig 3A) that interact only with the proteins (red particles in Fig 3A) in the PNC. This new species, which is the ligand, adsorbs to a protein with a rate *k_1_* and desorbs with rate *k_2_*, which together determine the ligand duty ratio 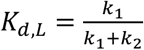. The ligand can also unbind from the membrane when the protein it is bound to desorbs from the actin aster. We have assumed that binding of the ligand to the membrane does not change its desorption rate, so that the ligand-protein complex desorbs with rate *k_off_*. Essentially, this model adds another layer of 1:1 Langmuir adsorption kinetics [30] on top of the protein-actin kinetics to understand the ligand-protein interactions on PNCs. Therefore, similar to the protein, the ligand can also form a cluster by adsorbing to the protein, which grows from *m* = 0 to *m* = *M_max_* at a propensity *k_1_n*, where *n* is the instantaneous number of proteins in the PNC. The ligand cluster decays at a propensity *k_2_m* and eventually returns to the *m* = 0 state, which again determines the desorption time of the ligand cluster. The ligand cluster lifetime is determined by three timescales: its own desorption time, the desorption time of the PNC, and the aster fragmentation time. The aster fragmentation time is unimportant for the desorption dominated PNCs, which allows us to infer the ligand lifetime solely from the ligand cluster and the PNC cluster desorption times.

**Figure 3.**
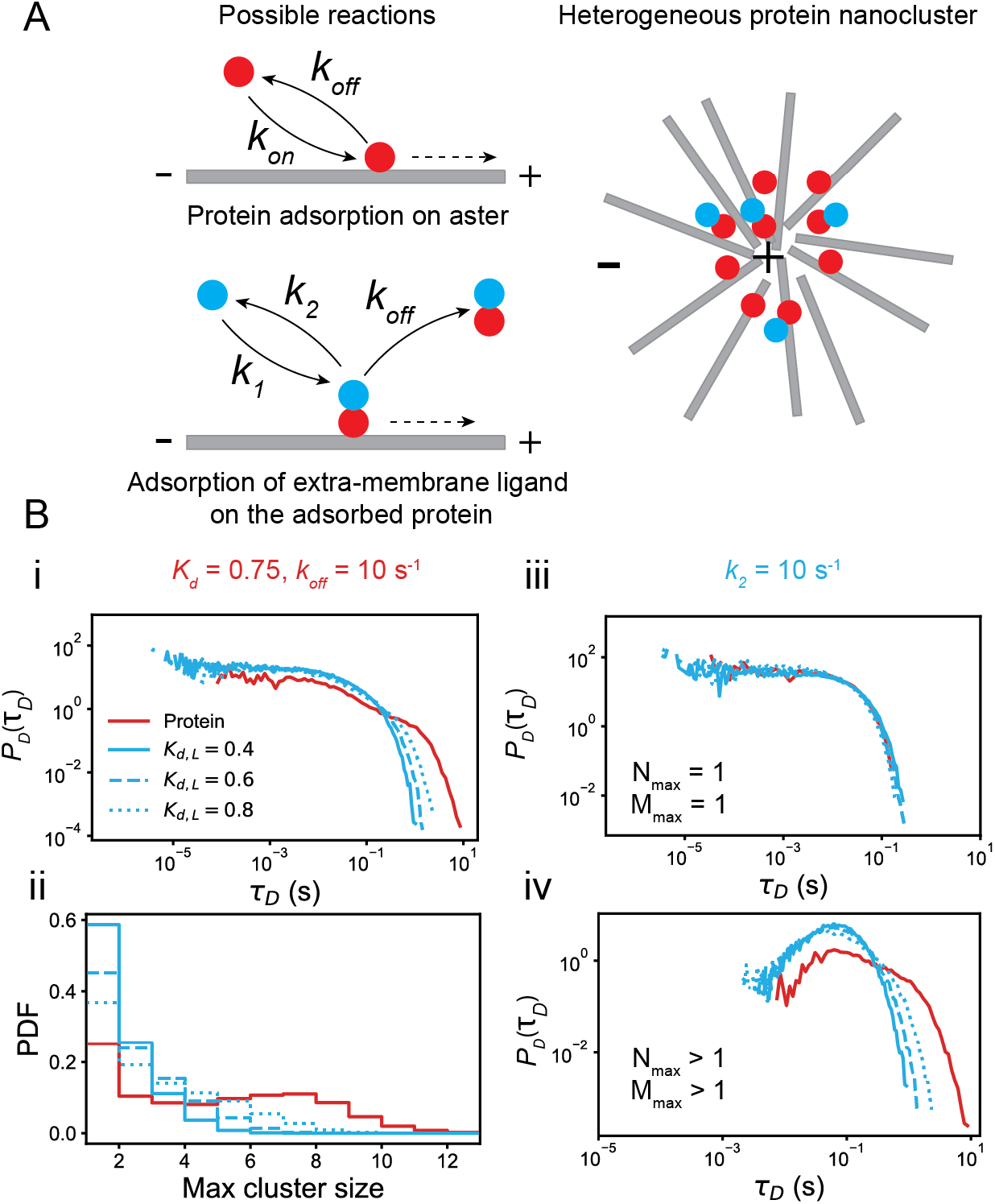
Ligand-Protein interaction on protein nanocluster. (A) The model from Fig.1(A) is amended to include interaction of the clustered protein target (red) with an extra-membrane ligand (blue). The ligand binds to the target with rate *k_1_* and dissociates with a rate k_2_. Therefore, in the presence of *n* targets, the number of the ligand, *m*, grows with propensity *k_1_* ×n and decays with propensity *k_2_* × *m*. In addition, the ligand-target complex also dissociate with propensity *k_off_* × *m*. Collectively, these reactions lead to the formation of a heterogeneous cluster, with sizes *N_max_* for the protein target (red) and *M_max_* for the ligand (blue). (B) Because *K_d_* = 0.75 and *k_off_* = 10s^−1^ fall in the desorption dominated region in Fig. 2B, the cluster lifetime is almost entirely determined by the desorption timescale *τ_D_*. In (i) we have plotted *P*(*τ_D_*) for both the target and the ligand and in (ii) we have plotted the cluster size distribution for various *K_d,L_* values (legend). As evident from (ii), the clusters with size 1 contribute heavily on *P*(*τ_D_*). To isolate the effect of cluster size, in (iii) we plot *P*(*τ_D_*) for *N_max_* = *M_max_* = 1, which decays exponentially with rate *k_off_*, whereas in (iv), *P*(*τ_D_*) for *N_max_,M_max_* > 1 have non-monotonic distributions. 10^4^ independent realizations of the model were used for these results.

The ligand-protein interaction varies widely depending on *K_d_,k_off_,K_d,L_*, and *k_2_*, some of which we have shown in Fig.S2. In the rest of the paper, we apply our general framework on the specific case of RAS-RAF interaction, which is a well-known model system. In particular, from biochemical measurements of the interaction between the Ras Binding Domain (RBD) of RAF and RAS, it has been shown that RAF dissociates from RAS approximately at a rate of *k_2_* ~ 10s^−1^ [31]. We also know that the average KRAS cluster contains about 6-8 proteins [3,4], which implies that in our model, *K_d_* ≈ 0.75 (Fig 2C (iii)). Finally, we also know that typical KRAS clusters survive for 0.1-1s [4], which we get when *k_off_*~10s^−1^ (Fig 2D). Therefore, *K_d,L_* is the only free parameter in our model. In Fig. 2B, cases with different *K_d,L_* is shown. Because binding of the ligand does not change the protein kinetics, the desorption time distribution of the PNC remains unaffected by changes in *K_d,L_*, but as *K_d,L_* increases, *k_1_* increases, which increases the propensity of ligand binding, and as a result the ligand cluster survives for longer as reflected in the broader tails. Due to the same reason, the size distribution of protein cluster remains unaffected, but the ligand cluster becomes larger as *K_d,L_* increases. Therefore, purportedly, the change in the ligand cluster lifetime distribution is due to the increase in the number of ligand clusters with size greater than one. Indeed, resolving the lifetime distribution by the size of the ligand and the protein clusters shows that the lifetime distribution of clusters of only one protein or ligand depends only on the desorption rates, whereas the lifetime of ligand clusters with *M_max_* > 1, changes with *K_d,L_*. Therefore, the formation of ligand clusters on PNCs provides *a* mechanism to control the residence time of extra-membrane ligands.

### RAF residence time on the membrane

Due to their roles in various cancers, understanding RAS-RAF interaction has been subject of extensive investigations, which has reported that RAS forms actin-dependent PNCs on the inner leaflet of the plasma membrane only when it is in the GTP-bound active form [2,4,32]. Also, it is well-known that RAF binds principally to GTP bound RAS [26]. Hence, it is likely that RAS-RAF interaction in human cells is mediated by RAS nanoclusters. Therefore, RAS-RAF interaction provides an excellent experimental platform for understanding the kinetics of ligand-cluster interaction on real PNCs.

To do so, we measured the residence times of RAS and RAF on Mouse Embryonic Fibroblast (MEF) cell membranes using TIRF microscopy and single-particle tracking (SI). To understand the effect of interaction between RAS and RAF on the residence time of RAF on the membrane, we used wild-type (wt) and mutated variants of RAS and RAF. The residence time of RAS decays as a power law with an exponential tail (Fig. 4A). This distribution is qualitatively similar to the distribution predicted by our model (Fig. 2A & 3B) except that our model predicts an exponential decay of the residence times. This is expected, because, in the model, we assumed that the actin concentrations are identical in all asters, which does not apply to a cellular system, where the cortical actin concentration can vary substantially over space and time. A simple way to incorporate this variation in our model is to assume that *k_on_* varies randomly from aster to aster due to spatial variation in *C*, but remains constant for an aster [33]. Indeed, if we assume that *k_on_* varies exponentially (Fig. 4E) with *k_off_* = 20*s*^−1^ and ⟨*K_d_*⟩ = 0.6, the *τ_C_* distribution from our model exactly reproduces the experimentally observed residence time distribution of RAS (Fig 4A & S3).

**Figure 4.**
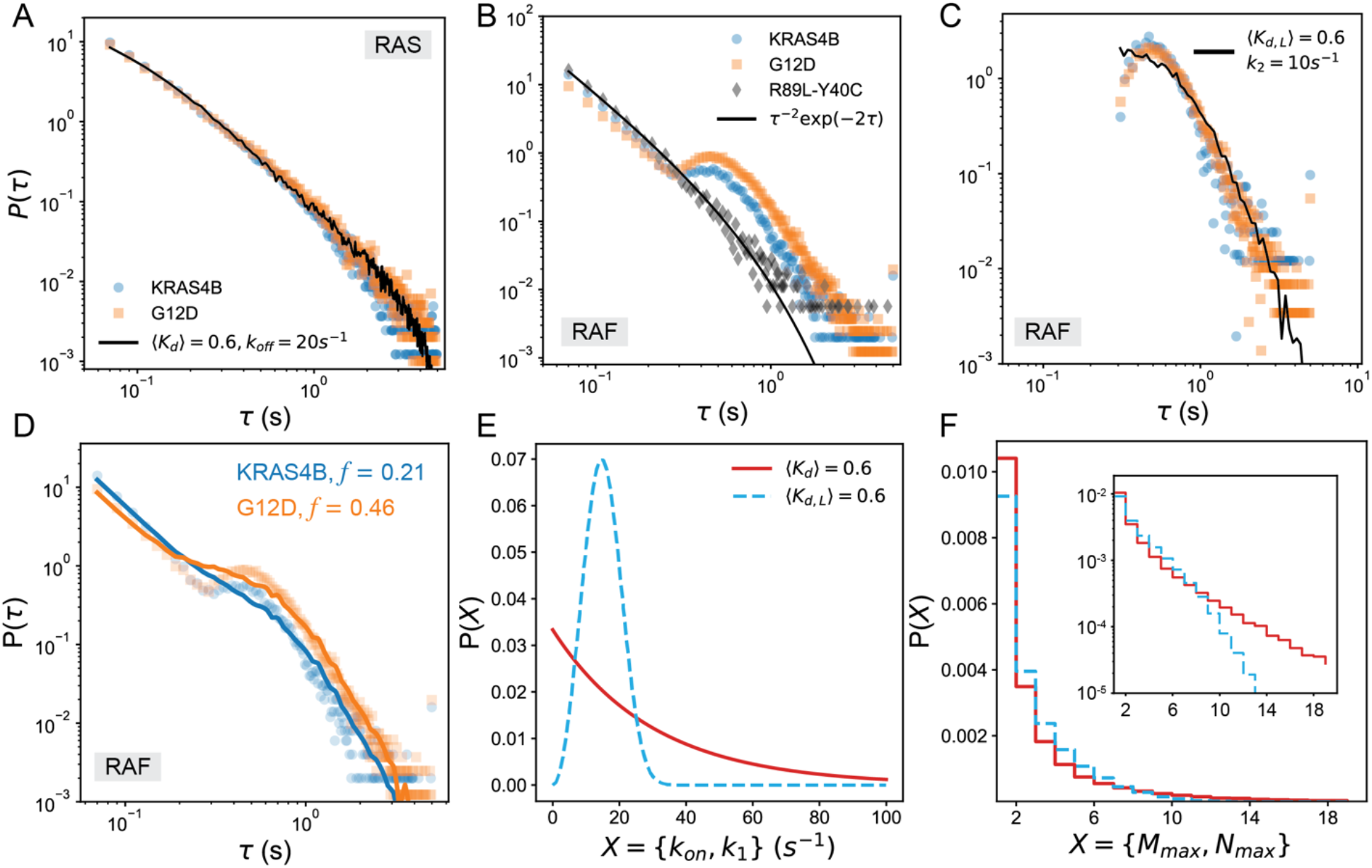
RAS-RAF interaction. (A) *RAS residence time:* Experimentally observed residence time distributions of wild type KRAS4B (blue circles) and KRAS4B with G12D mutation (orange square). The black line is the model prediction for a set of parameters (legend). (B-D) *RAF residence time:* (B) Experimentally observed residence time of c-RAF on the membrane in the presence of different RAS mutation (same color scheme as in (A)). Both distributions show power law decay at short times (*τ* < 0.25 *s*) and nonmonotonic decay at longer times. The power law is also observed in the presence of R89L mutation in RAS and Y40C mutation in RAF (black diamonds). These two mutations remove any interaction between RAS and RAF. Therefore, this observation suggests that the power-law for short times arises due to RAF-membrane interactions and the nonmonotonic decay at longer times arises due to RAS-RAF interactions. (C) Because these two processes are independent, we subtract the R89L-Y40C curve from the other two to get the RAF residence time distribution arising purely due to RAS-RAF interactions (blue and orange markers). We find good match with our theoretical predictions of *P*(*τ_c_*) (black curve) when two conditions are met: (1) *k_on_* (red in I) and *k_1_* (sky-blue I(E)) are randomly distributed, with ⟨*K_d_*⟩ = 0.6 and ⟨*K_d,L_*⟩ = = 0.6 values with k_off_ = 20s^−1^ & *k_2_* = 10s^−1^, and (2) we consider the distribution of residence time only when *M_mαx_* ≥ 8. (D) Adding our model’s prediction with the power law from membrane-RAF interaction reproduces (solid lines) experimental observations (markers) to an excellent deIe. (E) The distribution of random *k_on_* and *k_1_* values used in the model. (F) The cluster size distribution of RAS (red) and RAF (sky-blue) predicted from our model. *Inset:* The same distribution in semilog scale. 10^5^ independent realizations of the model were used to obtain each distribution.

The residence time of RAF on the membrane follows a nonexponential distribution (Fig 4B-D), which suggests that the residence time of RAF is *not* determined by uncorrelated collisions with the membrane and the interaction of RAF with RAS and the membrane are its important determinants. Indeed, upon closer inspection of the residence time distribution for wt.RAS and RAF (Fig. 4B) we found that the residence time, *τ*, has two unique regimes: for *τ* < 0.25 *s*, the distribution decays as a power law and above this timescale, it decays nonmonotonically (has a peak) with a different power law tail. To understand the origin of these two regimes, we repeated the experiments on RAS with R89L and RAF with Y40C mutations. The mutations eliminate any interaction between RAS and RAF. For this system, the nonmonotonic part disappeared from the residence time distribution and only the power law decay remained (Fig. 4B), which implies that the nonmonotonic part originates from the RAS-RAF interactions, whereas the initial power law decay originates from the RAF-membrane interaction. This is further evinced by the residence time distribution of RAF in the presence of RAS with G12D mutations, which increases the fraction of GTP bound RAS. In this system, the peak of the nonmonotonic part becomes more pronounced (Fig. 4B).

The above observations show that the initial power law and the nonmonotonic decay at later times originate from two independent processes. Hence, we can isolate the contribution of RAS-RAF interaction on the RAF residence time by subtracting the power law obtained from R89L-Y40C system from the other two residence time distributions. Doing so produces the distributions shown in Fig. 4C. The distributions are identical to each other within experimental variations and have power law tails that decay as τ^−3.5^. Remarkably, the shape of the distributions is qualitatively similar to the distribution of ligand cluster desorption times (Fig 3B). Hence, it is possible that the nonmonotonic distribution arises due to the formation of RAF clusters on the RAS clusters. However, it is also possible that the nonmonotonicity arises due to the multiple rebinding of a single RAF on the cluster, which increases its lifetime and leads to a nonexponential and nonmonotonic distribution. However, our computational results showed that although multiple rebinding of a single ligand increased the residence time, it did not make the residence time distribution nonmonotonic (Fig. S4), which ruled out multiple rebinding as a possible origin of the observed nonmonotonic distribution. Importantly, this result established that ligand clustering is the underlying mechanism of the nonmonotonic decay of residence times.

Similar to the RAS residence time distribution, the power law decay in RAF residence time distribution can be explained by the spatial variation of *k_1_* from one PNC to another because of the differences in local lipid environments or variation in effective reaction rates due to competition from other interaction partners of RAS. We incorporate the spatial variation by assuming that (besides *k_on_*) *k*_1_ is also randomly distributed (Fig. 4E). Remarkably, we find that introduction of the random *k_on_* and *k*_1_ suffices to reproduce the power law tail. In fact, we can quantitatively reproduce the distribution when we consider the lifetime of ligand clusters with *M_max_* ≥ *M_th_* = 8 and ⟨*K_d,L_*⟩ = 0.6 (Fig. 4C-D), which confirms that the nonmonotonicity arises solely because of the formation of RAF clusters on RAS clusters (Fig S5). Indeed, the cluster size distribution predicted by our model (Fig 4F) is consistent with prior experimental [4,19] and computational observations [19,34–36].

### Impact of the G12D mutation of RAS

The consistency of our model allows us to investigate the origin of the difference between the residence time distributions in the presence of wt and G12D RAS. In particular, *why do we see an increase in the nonmonotonic part of the distribution in the presence of G12D, even though the RAS-RAF binding affinity remains unchanged*? In our model, in the absence of any changes in the binding affinity, the nonmonotonic part can only get enhanced if a higher fraction of RAF became clustered through its interaction with RAS PNCs. To test this hypothesis, we added the experimentally obtained power law *P_E_*(*τ_E_*) (Fig 4B) to the cluster time distribution *P_C_(τ_C_*) from our model and generated a combined distribution by taking a weighted mean of the two distributions.

The resultant distribution is:

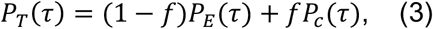

where *f* is the fraction of RAF that binds to RAS PNCs. We found a best fit distribution for both cases by varying the threshold *M_th_* (Fig. S5) and *f*. Remarkably, we found that both best fit distributions had identical *M_th_* = 8, but the *f* values were different by a factor of two (Fig. 4D). This result implies that, in the timescales probed by our experiments (~5 s), the G12D mutation does not change the binding affinity between RAS and RAF, but it increases the number of RAS clusters. This result is consistent with the experimental observation that only RAS.GTP forms clusters [4]. Because the G12D mutation effectively increases the number of RAS.GTP on the membrane, it is likely that we will observe more RAS clusters in the presence of this mutation.

## Discussion

In this paper, we have presented a simple mathematical model to understand the lifetime of actin-dependent peripheral membrane protein nanoclusters (PNCs) and protein-ligand interactions on the protein nanoclusters. Our results show that in many biologically relevant cases, the lifetime of PNCs is determined solely by the adsorption-desorption kinetics of the proteins on actin asters and not by the fragmentation of the aster. Our model shows that many PNCs arise from subcritical nucleation [37]of peripheral membrane proteins that survive for a short time before disintegrating. Under special circumstances, the nucleus becomes large enough to be stable and survives for a long duration. Only in these cases, the fragmentation of the aster determines the lifetime of the PNC. Therefore, from a physical standpoint, the dynamics of PNCs is better understood by studying the dynamics of subcritical nuclei, as we have done in this paper.

To understand the effect of protein clustering on the ligand-protein interactions, we studied some ideal cases using our model, which showed that the clustering of the ligands on the membrane enhances the residence time of the ligands on the membrane. We compared our results with experimental measurements of RAF residence times in the presence and absence of its interaction with the PNC forming RAS protein. We found remarkable agreement between our predictions and the experimental observations, which consolidated the results obtained from our model. Importantly, investigation of this system using our model allows us to contribute on an ongoing debate on the role of RAS G12D mutation on the proliferation of cancer cells. Our model suggests that the G12D mutation does not change the binding affinity between RAS and RAF. Instead, it increases the propensity of RAS and RAF cluster formation, which increases the residence time of RAF on the membrane and enhances the activity of the MAPK pathway involved in proliferation. An independent experiment on HeLa cells shows very similar results (not shown), which gives further credence to our proposition.

Although our model showed remarkable agreement between the experimental observations, there are several drawbacks that need to be addressed to develop a better model of the PNCs. We assumed well-mixed mass action kinetics, which is likely to be invalid in most biological context [38]. The well-mixed kinetics also overestimates the frequency of events happening at short times, because of which our model disagrees with the experimental observation at short times (Fig 4B-D). Also, in modeling the protein-ligand interactions, we did not explicitly model the multiple rebinding of a ligand on a PNC and assumed that it will be captured by some effective *k_on_* or *k_1_* values. Indeed, recalculation of the results by adding a well-mixed model of multiple rebinding does not change our results qualitatively (not shown). Also, the lipid environment of proteins plays an important part in determining the stability of the PNCs [4,9,34,39,40], which we do not consider here explicitly. In the future, we will develop models to incorporate these features. Despite these limitations, our current model provides deep insights into the working of the PNC formation and lifetime which will be useful in our understanding of protein nanoclusters and protein-protein interactions during cell signaling.

In conclusion, in this paper, we present a general framework to study protein nanocluster dynamics. As demonstrated here, through this framework, we can not only study general questions regarding the growth and stability of protein nanoclusters, but also apply it to study specific protein-protein interaction systems. The framework presented is not specific to RAS-RAF system and can be used to model other protein-protein interactions just by varying the parameters. Furthermore, with little modification, our framework can be used to understand drug-protein interactions, which may be useful in rapid design of novel drugs. We believe the generality and the simplicity of our framework will be useful in studying various biomolecular interactions and provide novel insights into their dynamics.

## Materials & Methods

Experimental techniques and theoretical methods are described in the SI.

## Acknowledgement

The authors would like to thank Angel E. Garcia, Dwight V. Nissley, Sumit K. Majumder and Van A. Ngo for critical reading of the manuscript. SS acknowledges funding support from an LDRD grant (XX01) from LANL. This work has the following LA-UR number: LA-UR-21-30526. This project has also been funded in whole or in part with Federal funds from the National Cancer Institute, National Institutes of Health, under Contract No. HHSN261201800001I. The content of this publication does not necessarily reflect the views or policies of the Department of Health and Human Services, nor does mention of trade names, commercial products, or organizations imply endorsement by the U.S. Government.

