## Supplementary Information for "Lifetime of actin-dependent protein nanoclusters"

#### Experimental materials & methods

**Cell Culture, transfection and labeling of HaloTag-Ras**

Mouse embryonic fibroblast (MEF) cells were genetically modified to express SNAP Tag fusion RAF and HaloTag fusion RAS protein. Fusion constructs were incorporated in cells using viral delivery methods and selected with antibiotics for stable expression. MEF cells plated and grown without antibiotics for 48 hrs before imaging. For the experiments with over-expression system, HeLa cells were transfected with HaloTag fusion constructs of various RAF isoforms in six well plates. Transfection was conducted using Fugene (Promega) reagent and 1.5ug DNA per well. The day after transfection, cells were transferred on to clean coverglass (#1.5, plasma-cleaned) and allowed to grow for another day. On the day of imaging, coverslips were washed with phosphate buffer saline and cells were labelled with 100uM fluorescent (JF646 or JF549) HaloTag ligand and SiR647, which covalently binds to the HaloTag-RAS molecules and SNAP Tag-RAF molecules respectively. Fluorescent labeled HaloTag ligands were obtained from Dr. Luke Lavis at (HHMI, Janelia Farm, Ashburn, VA). These fluorescence dyes are highly photostable and resistant to photobleaching (Grimm, 2016 #16).

**Single molecule microscopy**

Single molecule imaging was carried out on the Nikon N-Storm microscope equipped with an APO x100 TIRF objective of 1.49NA (Nikon, Japan). A Tokai hit stage incubator (Tokai Hit Co., Ltd, Japan) was used to provide 5% $CO_2$ while maintain the temperature at 37° for live cells. Labeled molecules (with JF646/JF549 or SiR647 dyes) associated with membrane were illuminated under TIRF mode. The JF549 dye was excited with the 561nm laser which is one of the four laser lines from the Agilent laser module of the Nikon N-STORM system (Serge et al., 2008), the JF646 or SiR647 dyes were excited with the 647nm laser line. The output laser beam was coupled into the Nikon TIRF box through a single mode fiber and focused into the back focal plane of the objective to form a parallel beam for wide field operation. The TIRF illumination was achieved by changing the illumination angle through the Nikon TIRF box controlled by the Nikon software (NIS- Elements AR 4.4). Fluorescent signals from each molecule were recorded with a thermoelectric-cooled EM-CCD camera with 16µm pixel size, (iXon Ultra DU-897, Andor Technologies, USA). Single molecule tracking was implemented by time-lapse imaging of the molecules under continuous illumination at 10ms exposure for a total of up to 2000 frames with zero delay time between frames. At this frame rate, membrane bound molecules appear as transient, diffraction-limited fluorescence spots. An area of 16x16 $\mu m^2$ of the plasma membrane in the cytoplasmic region of each cell was imaged.

**Single molecule tracking data processing**

The ImageJ-based single molecule tracking plugin, TrackMate [1] was used to create tracks from the time-lapse movies. Single molecules were identified as spots from each frame of the time-lapse movies with the eight-way adjacency particle detection algorithm with 30 GLRT [2] sensitivity and a PSF of 1.3 pixels. Sub-resolution spot accuracy was achieved using a 2D Gaussian fit function for estimating the position of the PSF for each frame. These spots were linked into tracks given certain criteria and cut off. The single molecule spot detection and tracking parameters were kept consistent across all experiments. These tracks were organized and exported for residence time analysis using a semi-automated workflow, developed in Matlab

(Mathwork, Natick, MA), on a multi-core Mac Pro. Tracking data was obtained in multiple replicates for each and every condition (~10000 tracks and 20 cells). Residence time was calculated from each track using a custom-developed Matlab routine.

### Theoretical Methods

#### Mathematical model of protein nanoclusters

The Langmuir kinetics [3] of protein adsorption-desorption can be summarized through the following reactions:

1. Adsorption of a protein and assimilation to the cluster of size $n$, denoted $P_n$ with rate $k_{on}$. The propensity of adsorption is $k_{on}$.
$$P_n \xrightarrow{k_{on}} P_{n+1}$$
2. Desorption of a protein from a cluster of size $n$ with rate $k_{off}$ and propensity $k_{off}n$.
$$P_n \xrightarrow{k_{off}} P_{n-1}$$

#### Mathematical model of ligand-protein interaction

1. Adsorption of a ligand to a protein with rate $k_1$, which for a cluster of size $n$, happens with intensity $k_1 n$. The adsorption of a ligand changes the size of the ligand cluster of size $m$, $L_m$, by one.
$$L_m P_n \xrightarrow{k_1} L_{m+1} P_n$$
2. A ligand desorbs from a ligand cluster of size $m$ with rate $k_2$ and propensity $k_2 m$.
$$L_m P_n \xrightarrow{k_2} L_{m-1} P_n$$
3. A ligand-protein complex desorbs with rate $k_{off}$ and propensity $k_{off}n$ to reduce both the ligand and the protein cluster size by one.
$$L_m P_n \xrightarrow{k_{off}} L_{m-1} P_{n-1}$$

#### Solution of the mathematical models

The number of proteins in a cluster is very small (usually $< 30$), hence the kinetics becomes nondeterministic due to intrinsic noise originating from small copy number of the proteins. As a result, we cannot use ordinary differential equation (ODE) based chemical kinetic models and have to solve these models using chemical master equations (CME). The CME for the reactions described here were solved using the Gillespie algorithm [4]. To generate the distributions in Fig 2 and 3, for each set of parameters, $10^4$ independent time series were generated by solving the model using the Gillespie SSA. Each of these time series were run for a maximum of 1000s or until the proteins completely desorbed from cluster, such that the cluster size became 0, whichever was shorter.

#### Distribution of $\tau_C$

The cluster lifetime $\tau_C$ is given by the minimum of the aster fragmentation time $\tau_A$ and the protein cluster desorption time $\tau_D$. That is $\tau_C = \min(\tau_A, \tau_D)$. We assume that the probability density distributions $P_A(\tau_A)$ and $P_D(\tau_D)$ are nonidentical, but independent of each other because the aster fragmentation and the protein desorption are independent processes for the clustering proteins that we consider here. They won't be independent if the clustering of the proteins, e.g., Myosin or Arp2/3 complex, directly influences the formation of the asters.

To find the distribution of $\tau_C$, $P_C(\tau_C)$, we use the cumulative distribution function (CDF) trick. To perform this trick, observe that:

*Obs 1:* $\tau_C > \tau$ *if and only if* $\tau_D > \tau$ *and* $\tau_A > \tau$.

Therefore,

$$P_c(\tau_C > \tau) = P(\tau_D > \tau \; and \; \tau_A > \tau)$$

Using the independence of the probability distributions of $\tau_A$ and $\tau_D$, and using the fact that the CDF $F_c(\tau) = 1 - P_C(\tau_C > \tau)$, we can rewrite the above expression as:

$$F_c(\tau) = 1 - P_D(\tau_D > \tau)P_A(\tau_A > \tau) = 1 - [1 - F_D(\tau)][1 - F_A(\tau)]$$

The distribution $P_c(\tau) = \frac{\partial F_c(\tau)}{\partial \tau}$. Therefore, the distribution is given by:

$$P_c(\tau) = S_A(\tau)P_D(\tau) + S_D(\tau)P_A(\tau),$$

where $S_X(\tau) = 1 - F_X(\tau)$ is the *survival function.*

**Measurement of overlap**

We measured the overlap between two probability distributions using the Bhattacharya coefficient ($BC$) [5]. The Bhattacharya distance $D_B$ (defined below) and the Kullback-Leibler divergence ($D_{KL}$) [6], which is usually used to measure the distance between two probability distributions give quantitatively similar results. The advantage of the Bhattacharya distance is that it is symmetrical for both distributions. For two continuous probability density functions $P$ and $Q$, these measures are defined as follows:

$$BC(P, Q) = \int_{-\infty}^{\infty} \sqrt{P(x)Q(x)} dx$$
$$D_B = -\ln BC(P, Q)$$
$$D_{KL}(P||Q) = \int_{-\infty}^{\infty} P(x) \log\left(\frac{P(x)}{Q(x)}\right) dx$$

**Best fit distribution**

To find the best fit distributions shown in Fig 4C and Fig 4D, we minimized the distance between the experimentally observed distribution and the theoretical distribution obtained from the weighted sum of $P_E(\tau_E)$ and $P_C(\tau_C)$:

$$P_T(\tau) = f P_C(\tau) + (1 - f)P_E(\tau)$$

The distances were measured using the Bhattacharya distance $D_B$ and the KL divergence $D_{KL}$, both of which predicted identical values of $M_{th}$ and $f$.

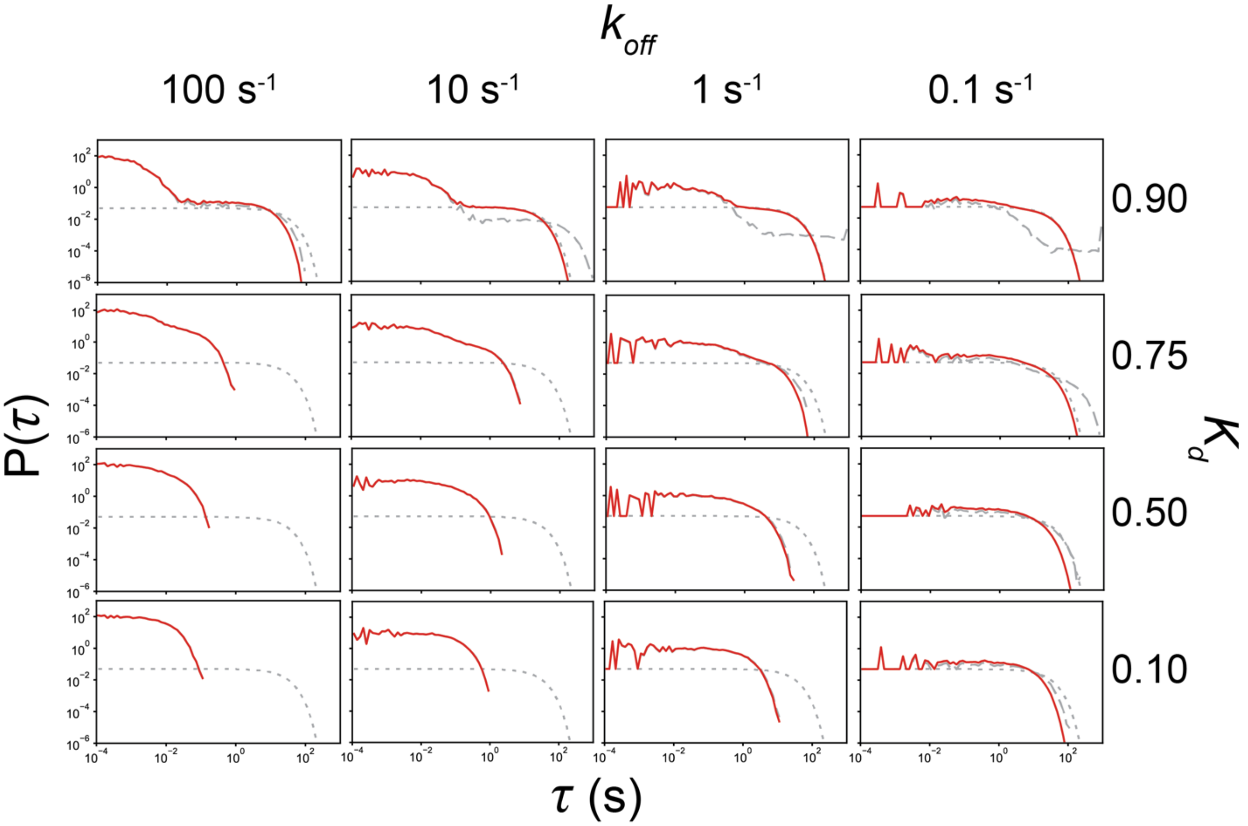


**Figure S1.** The lifetime distribution of protein adsorbed on actin aster. The aster lifetime $\tau_A$ has exponential distribution, $P_A(\tau_A)$with mean lifetime of $20\ s$ (grey dotted line). The desorption lifetime distribution $P_D(\tau_D)$ is shown using a gray dashed line and the cluster lifetime distribution, $P_C(\tau_C)$ is shown in red.

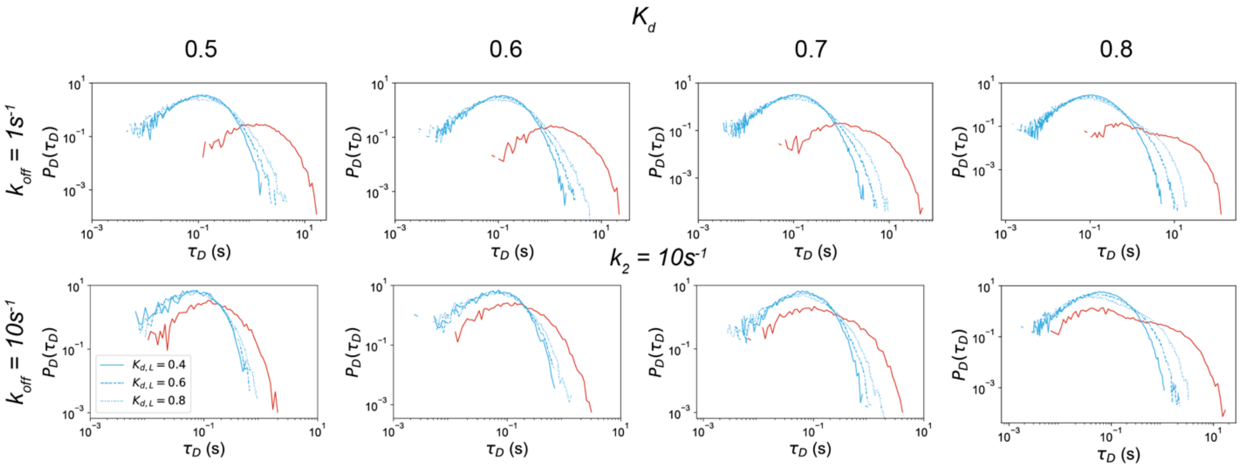


**Figure S2.** Desorption time distribution $P_D(\tau_D)$ of proteins (red) and ligands (sky-blue) when $N_{max} \geq 2$ and $M_{max} \geq 2$. The parameter values are shown in the figure.

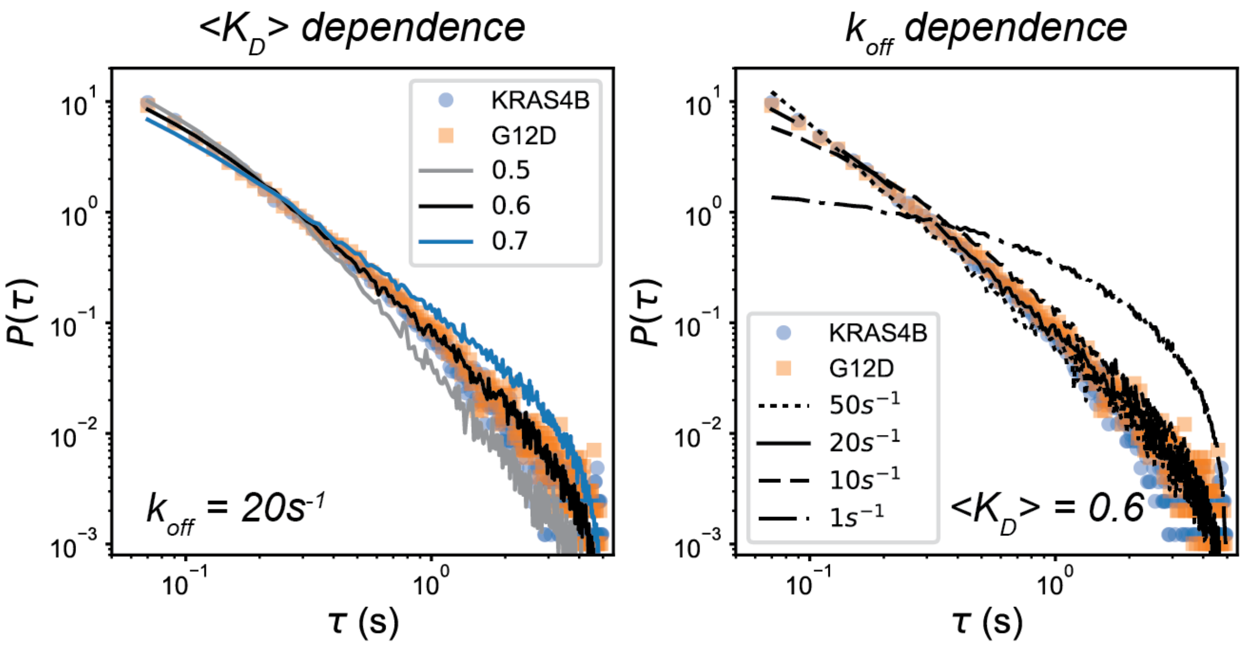


**Fig S3.** Fitting the model predictions of PNC lifetime distribution for various values of $\langle K_d \rangle$ for $k_{off} = 20\ s^{-1}$ (left) and $k_{off}$ for $\langle K_D \rangle$ (right), which shows that the theoretical curves best fit the experimental data when $\langle K_D \rangle = 0.6$ and $k_{off} = 20s^{-1}$.

**Rebinding of a single ligand: a chemical kinetic model**

A ligand molecule may rebind to a protein target after dissociating from the target. The rebinding of the ligand enhances its lifetime near the protein. This effect is enhanced near protein nanoclusters, where the ligand can find a large number of targets to rebind before diffusing away into the bulk. Therefore, the residence time distribution of a ligand can change drastically near a protein nanocluster.

To understand the effect of rebinding on the residence time distribution of a single ligand molecule, we consider the following reactions:

1. Free ligand entering a region (reaction domain) where it can interact with the proteins in the nanocluster.
$$\phi \xrightarrow{k_{cm}} F$$
2. Free ligand diffusing into the bulk.
$$F \xrightarrow{k_{mc}} \phi$$
3. Free ligand interacting with the protein target to form ligand-protein complex.
$$F + P \overset{k_1}{\rightarrow} PR$$
4. Ligand-protein complex dissociating to produce a free ligand.
$$PR \overset{k_2}{\rightarrow} F + P$$
5. Ligand-protein complex dissociating from the membrane and diffusing to the bulk.
$$PR \xrightarrow{k_{off}} \phi$$

In addition, the protein nanocluster dynamics is governed by the reactions described in Fig 1A. To ensure that there is only one ligand in the reaction domain, reactions 1 and 4 can occur only when $F = 0$ and reactions 2 and 3 can only occur when $F = 1$. As a result, the ligand can bind to / unbind from the protein multiple times before escaping to the bulk.

We estimate $k_{cm}$ by finding the diffusion limited rates of binding to an absorbing spherical domain of radius $a$, which is the radius of the reaction domain. Because the reaction domain is hemispherical, $k_{cm}$ will be proportional to the value for a spherical domain. The absorption rate of a spherical domain depends on the bulk concentrations, $c_\infty$, of the ligands and their diffusivities, $D$. For purely diffusive transport, $k_{cm}$ will be proportional to $4\pi D a c_\infty$, where $a$ is the radius of the hemispherical reaction domain [7]. For $c_\infty = 200/\mu m^3$, $D = 10\ \mu m^2/s$, and $a \approx 5 - 50\ nm = 0.005 - 0.05\ \mu m$ which are the typical numbers for signaling protein clusters and their protein partners, $k \approx 100 - 1000/s$.

Similarly, the escape rate $k_{mc}$ can be approximated from the first escape time, $\tau$, from a spherical domain of radius $a$, for which the escape time distribution is given by [8,9]:

$$q(\tau) = -2 \sum_{n=1}^{\infty} (-1)^n \exp\left(-\frac{n^2\pi^2}{a^2} Dt\right) \frac{n^2\pi^2 D}{a^2}$$

From this distribution, it is easy to compute the mean escape time, which for the same values of $D$ and $a$ is around $10^{-5} - 10^{-4} s$, which implies that $k_{mc} \approx 10^4 - 10^5/s$. The residence time distribution of the ligand for these values of $k_{mc}$ and $k_{cm}$ are shown in the following figure. Clearly, the residence time does not show any nonmonotonic behavior.

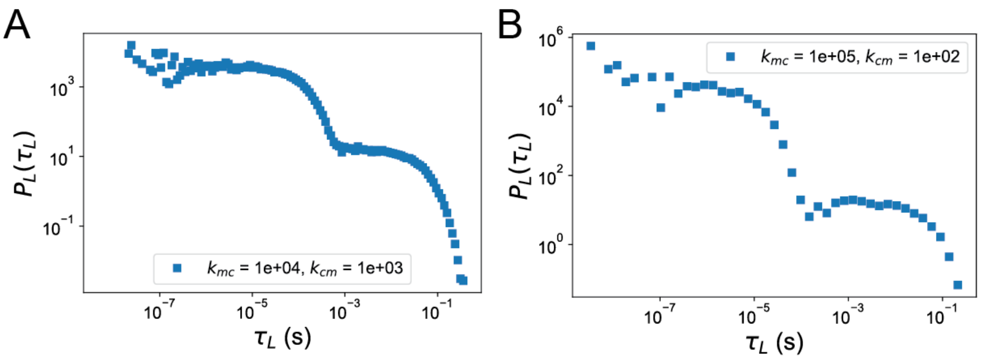


**Fig S4**. Residence time distribution of single ligand for biologically relevant parameter values does not show nonmonotonic distributions. $\langle K_d \rangle = 0.6$, $k_{off} = 20s^{-1}$, $\langle K_{d,L} \rangle = 0.6$, $k_2 = 10s^{-1}$. $k_{on}$ is distributed as an exponential and $k_1$ is distributed as a Weibull distribution. In both cases the mean values are determined by the mean duty ratios and the dissociation rates.

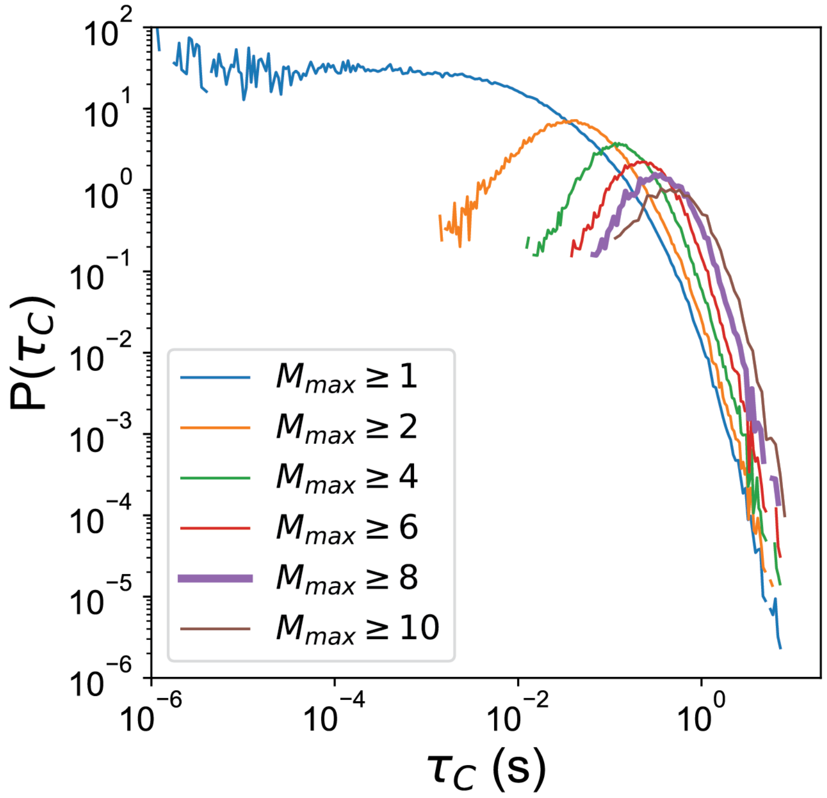


**Figure S5.** The cluster lifetime distribution $P(\tau_C)$ of the ligand (RAF) when $k_1$ is distributed as a Weibull distribution with shape parameter 3 and mean determined by $\langle K_{d,L} \rangle$ = 0.6 and $k_2 = 10s^{-1}$. We plot the distributions of $\tau_C$ for $M_{max} \geq M_{th}$. The values of $M_{th}$ are shown in the legend. For this plot $k_{off} = 20s^{-1}$, $\langle K_d \rangle = 0.6$, and that $k_{on}$ is distributed as a Weibull distribution with shape parameter 1 (exponential distribution).
